# Increased mitochondrial Ca^2+^ contributes to health decline with age and Duchene muscular dystrophy in *C. elegans*

**DOI:** 10.1101/2022.07.08.499319

**Authors:** Atsushi Higashitani, Mika Teranishi, Yui Nakagawa, Yukou Itoh, Surabhi Sudevan, Nathaniel J Szewczyk, Yukihiko Kubota, Takaaki Abe, Takeshi Kobayashi

## Abstract

Sarcopenia is a geriatric syndrome characterized by an age-related decline in skeletal muscle mass and strength. Here, we show that suppression of mitochondrial calcium uniporter (MCU)-mediated Ca^2+^ influx into mitochondria in the body wall muscles of the nematode *Caenorhabditis elegans* improved the sarcopenic phenotypes, blunting movement and mitochondrial structural and functional decline with age. We found that normally aged muscle cells exhibited elevated resting mitochondrial Ca^2+^ levels and increased mitophagy to eliminate damaged mitochondria. Similar to aging muscle, we found that suppressing MCU function in muscular dystrophy improved movement via reducing elevated resting mitochondrial Ca^2+^ levels. Taken together, our results reveal that elevated resting mitochondrial Ca^2+^ levels contribute to muscle decline with age and in muscular dystrophy. Further, modulation of MCU activity may act as a potent pharmacological target in various conditions involving muscle loss.

## Introduction

Sarcopenia is an age-related skeletal muscle disorder characterized by the accelerated loss of muscle mass and strength^1,2,3^. The skeletal muscles of aged mice and older people both display mitochondria with altered features including decreased volume, irregular morphology, and decreased functional activities^4,5,6^. Mitochondrial Ca^2+^ has been shown to regulate crucial mitochondrial functions such as energy production, reactive oxygen species (ROS) production, and the initiation of cell death^7^. Recently, it has been reported that aged mice display significantly increased levels of resting mitochondrial Ca^2+^ in skeletal muscle fibers^4^, indicating that dysregulated mitochondrial Ca^2+^ homeostasis could be involved in sarcopenia. Mitochondrial calcium uniporter (MCU) has been identified to be the primary channel responsible for mitochondrial Ca^2+^ uptake across the inner mitochondrial membrane^8,9^. Therefore, suppression of MCU function was expected to prevent undesirable accumulation of Ca^2+^ in mitochondria and to ameliorate sarcopenia. However, Rizzuto’s group has demonstrated that MCU silencing in rodent skeletal muscle causes muscle atrophy, and that overexpression of MCU in the hindlimb of mice, increasing mitochondrial Ca^2+^ uptake, causes muscle hypertrophy and provides a protective effect against denervation-induced atrophy^10^. Conversely, mutations in an MCU regulator, MICU1, which increase resting mitochondrial Ca^2+^ levels, caused neuromuscular disorders with cognitive decline, muscle weakness, and an extrapyramidal motor disorder^11^. Discrepancies in these findings indicates multiple roles of mitochondrial Ca^2+^ on skeletal muscle homeostasis. Namely, mitochondrial Ca^2+^ may have distinct effects on various processes of skeletal muscles aging. Therefore, to resolve the role of mitochondrial Ca^2+^ in sarcopenia, it is crucial to verify individually its role in a simple or prebiotic experimental system.

The body wall muscle in *Caenorhabditis elegans* has a structure similar to vertebrate skeletal muscle containing sarcomeres. Also similar to mammalian muscle, *C. elegans* muscle displays structural and functional declines with age^12,13^. *C. elegans* sarcomeres and mitochondria are located in the monolayer within the cell and can be easily observed alive under a microscope. Declines in mitochondrial network structure, increased fragmentation, and reduced mitochondrial volume occur earlier than sarcomere decline and correlate more strongly with reduction in movement, maximum velocity, and lifespan^14,15,16^. In addition, several molecular systems such as the dystrophin complex and mitophagy, which is controlled by PINK and PERKIN, are conserved in *C. elegans*^17,18,19^. Furthermore muscle deterioration can be examined without the influence of muscle regeneration since *C. elegans* has no muscle stem cells. Therefore, the *C. elegans* body wall muscle is a simple model useful to study (primary) sarcopenia and other inherited muscular diseases.

In this study, we examined the role of mitochondrial Ca^2+^ homeostasis in the context of aging and sarcopenia using *C. elegans*. Initially, like past studies, we observed aberrant changes in mitochondria in the body wall muscle of aged worms. We next confirmed elevated levels in resting mitochondrial Ca^2+^ with age. Either pharmacologic or genetic inhibition of MCU function was sufficient to prevent increases in mitochondrial Ca^2+^ and improve sarcopenic phenotypes. In addition, we found that Duchene muscular dystrophy (DMD) worms also exhibit abnormally high cytosolic and mitochondrial Ca^2+^ levels, and that MCU inhibition was similarly sufficient to prevent increases in mitochondrial Ca^2+^ and improve health. The results in this study indicate that altered mitochondrial Ca^2+^ homeostasis is associated with muscle aging and dystrophy in *C. elegans*. These findings raise the possibility that mitochondrial Ca^2+^ homeostasis is associated with mammalian muscle aging and dystrophy and that it may be a potential therapeutic target in them.

## Results and discussion

### Age-related mitochondrial changes with Ca^2+^ accumulation

In the body wall muscle on D4 adulthood of WT worms, the mitochondria contained aligned filamentous structures, but these structures fragmented and shortened with increasing age (Figures 1A and B). Our qualitative analysis showed that more than 70% of muscular cells in the D10 WT worms were classified into ‘fragmented’ or ‘very fragmented’. Next, we examined whether resting mitochondrial Ca^2+^ levels were elevated in the muscle cells of aged nematodes as previously observed in rodent models^4^. To estimate mitochondrial Ca^2+^ concentrations, we used genetically modified WT worms that have mitochondria-targeted red fluorescent Ca^2+^ indicator (mito-LAR-GECO1.2 (mtGECO))^20^ (*aceIs1* transgene; see Supplementary Figure S1A) and mitochondrial-and nuclear-targeted GFP (mtGFP and nucGFP) in their muscle cells (*ccIs4251* transgene)^21^. We imaged fluorescent signals of mtGECO in mtGFP-positive structures and calculated the mtGECO/mtGFP ratio. In the presence of ionomycin and Ca^2+^, mtGFP-positive structures in muscle cells of aged worms (D10) had about the same mtGECO/mtGFP values as D1 (Supplementary Figure S2A and B). However, in an intact state, mtGECO fluorescence on the mtGFP-positive structures of aged worms was observed to be more intense and the mtGECO/mtGFP values were found to be larger in aged worms (Figures 1A; Supplementary Figure S2C and D). Since the *Kd* of mtGECO (LAR-GECO1.2) is relative high (12 μM)^20^, there were thought to be age-dependent micromolar changes in mitochondrial Ca^2+^ concentration ([Ca^2+^]_mito_). Our quantitative analysis of [Ca^2+^]_mito_ estimated an increase with age from 0.6 ± 0.3 μM (D4) to 3.4 ± 2.8 μM (D10), and 5.6 ± 4.6 μM (D13) (Figure 1A and C), consistent with prediction. This is also consistent with results from a previous report using rodents, where mitochondrial Ca^2+^ levels increased with age^4^.

**Figure 1.**
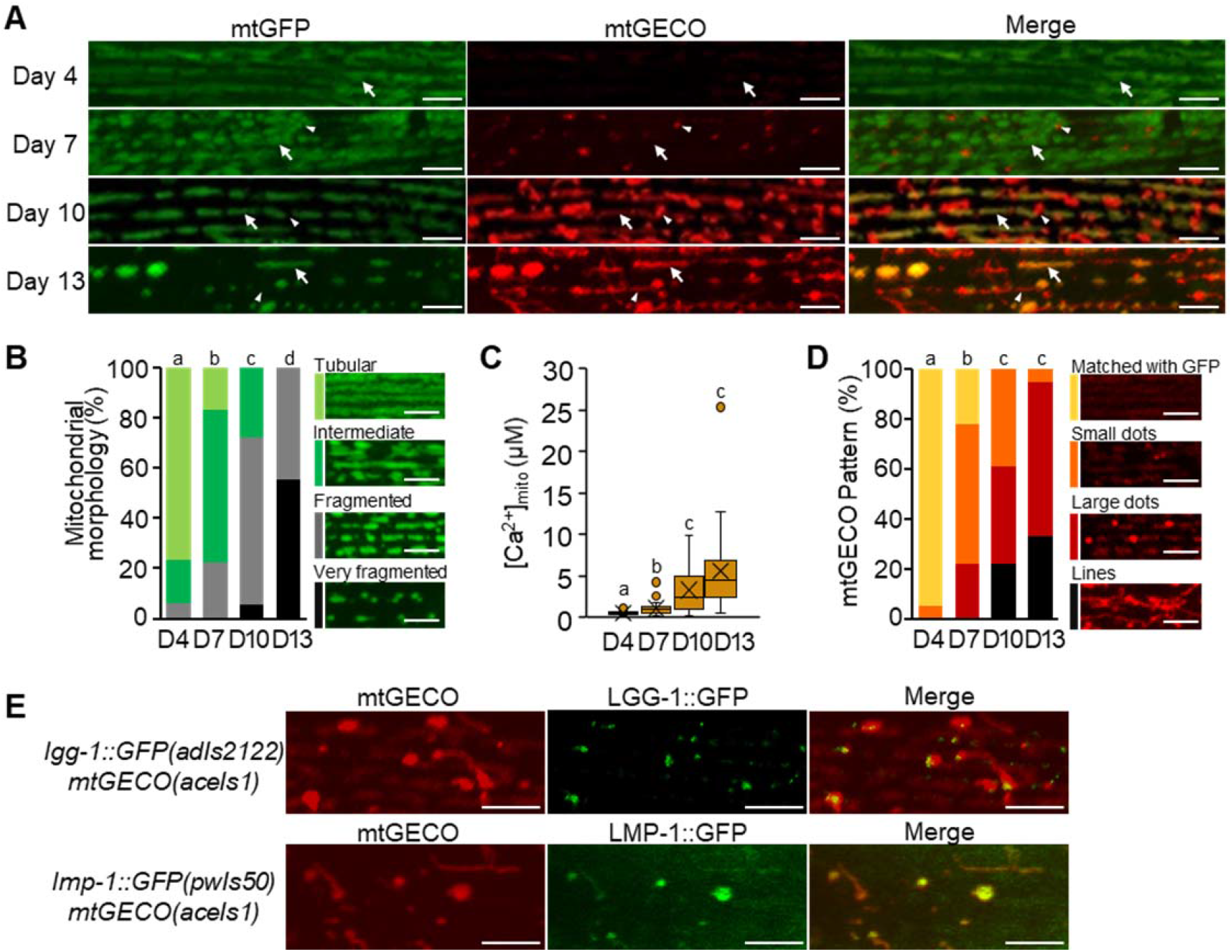
Decreasing in mitochondrial volume and increasing in mitochondrial Ca^2+^ level with age. (A) Representative fluorescence images of body wall muscle cells in transgenic *C. elegans* expressing mitochondria-targeted GFP (mtGFP) and a mitochondrial Ca^2+^ probe (mtGECO) in the body wall muscle cells in WT worms (*ccIs4251; aceIs1*) at D4, D7, D10 and D13 of adulthood. The arrows indicate the typical position where Ca^2+^ concentration in mitochondria ([Ca^2+^]_mito_) was measured in (C). The arrowheads indicate the position of mtGECO structures that no longer colocalized with mtGFP. Scale bars, 5 μm. (B) Qualitative analysis of body wall muscle cells with abnormal mitochondrial morphology. On the left, light green, green, light grey and black represent fractions containing cells with ‘tubular’, ‘intermediate’, ‘fragmented’ and ‘very fragmented’ mitochondria, respectively. The right panels show representative images of the categorized mitochondria. Scale bars, 5 μm. Data represent the value from 18 cells from at least 6 worms. (C) Age-dependent elevation in [Ca^2+^]_mito_ of muscle cells. Calcium levels of mtGFP-positive mitochondria as shown by arrows in (A), were calculated from the ratio of fluorescence intensity of mtGECO to that of GFP in a constant area described in Methods and Protocols. Data represent the value from 19-21 cells from at least 6 worms. (D) Qualitative analysis of body wall muscle cells with different mtGECO pattern. On the left, yellow, orange, red and black represent fractions containing cells with the mtGECO signal ‘matched with GFP’, and the fractions of the cells that showed ‘small dots’, ‘large dots’ and ‘lines’ of mtGECO signal, respectively. The right panels show representative images of the categorized patterns. Scale bars, 5 μm. Data represent the value from 18 cells from at least 6 worms. (E) Colocalization of mitophagy-related proteins with mtGECO-accumulated dots and tubular structures in D10 muscle cells. Representative fluorescence images of transgenic *C. elegans* (*adIs2122* [*Plgg-1*::*lgg-1*::*GFP*]; *aceIs1*) expressing mtGECO and GFP-tagged LGG-1 and transgenic *C. elegans* (*pwIs50* [*Plmp-1*::*lmp-1*::*GFP*]; *aceIs1*) expressing mtGECO and GFP-tagged LMP-1. Scale bars, 5 μm. Letters on the tops of bars indicate statistical significance by the chi-square test (B, D) or one-way ANOVA with Dunn’s multiple comparison test (C) (*p* < 0.05).

In *C. elegans*, age-related mitochondrial fragmentation and disappearance have previously been reported^14,15,16^. What is new in our study is that we found increased intramitochondrial Ca^2+^ levels (Figure 1A and C). Interestingly, we also found a greater extent in the mtGECO structures (‘small dots’, ‘large dots’ and ‘lines’) that no longer colocalized with mtGFP (Figure 1A and D). The high-intensity dots of mtGECO frequently appeared adjacent or connected to mtGFP structures. We suspected that the high-intensity dots of mtGECO were Ca^2+^-accumulated portions of mitochondria that would be eliminated by mitophagy to clear damaged mitochondria with age. This may be caused by RFP fluorescence of mtGECO persisting, while the mitochondrial GFP fluorescence quenched. Indeed, 20 minutes of live imaging confirmed that the mtGECO-only structures were isolated from the mitochondrial network, where both mtGFP and mtGECO were positve (Supporting Movie 1). In addition, the mtGECO structures colocalized with the autophagosomal marker LGG-1::GFP, or with the lysosomal marker LMP-1::GFP in D10 muscle cells (Figure 1E). Thus, mtGFP-negative mtGECO structures were trafficked into acidic lysosomal compartments, which must have quenched the fluorescence of GFP. In a weakly acidic environment, mtGECO can fluoresce but decrease fluorescence emission as the pH decreases^20^. In addition, mtGECO can have Ca^2+^-dependent fluorescence changes under a weak acidic condition (pH 5.5) judging from^20^. Therefore, we assumed that intense fluorescence intensities observed in the mtGECO structures could reflect Ca^2+^ accumulation. Taken together, these results suggest that the fragmented mitochondrial network in aged *C. elegans* muscle cells contain fragments with elevated Ca^2+^ levels, and that these Ca^2+^-accumulated portions of mitochondria are eliminated by mitophagic pathways, presumably to maintain mitochondrial function.

### Effect of loss-of-function mutation in *mcu-1* on muscle aging in *C. elegans*

ATU2301 (WT) adult *C. elegans* expressing the genetically encoded Ca^2+^ indicator GCaMP in muscle cytosol (*goeIs3*)^22^ and mitochondrial Ca^2+^ indicator mtGECO in muscle mitochondria (*aceIs1*) were immobilized on polystyrene microspheres and observed live over time. In these worms, cytosolic and mitochondrial Ca^2+^ levels fluctuated in synchronization with contraction and relaxation of the body wall muscles (Figures 2A, Supporting Movie 2). When a null mutation in the mitochondrial calcium uniporter *mcu-1*(*ju1154*) was introduced into this recombinant (ATU2302; *mcu-1*(*ju1154*)), cytosolic Ca^2+^ changes associated with muscle contraction were observed, but synchronized Ca^2+^ influx into the mitochondria was largely lost (Figure 2A, Supporting Movie 2). These results indicate that when a large amount of Ca^2+^ flows into the muscle cytosol due to muscle contraction, Ca^2+^ is also taken up in mitochondria via MCU-1. Gibhardt et al (2016) report similar usefulness of LAR-GECO as a mitochondrial Ca^2+^ sensor in the mammalian muscular system^23^.

**Figure 2.**
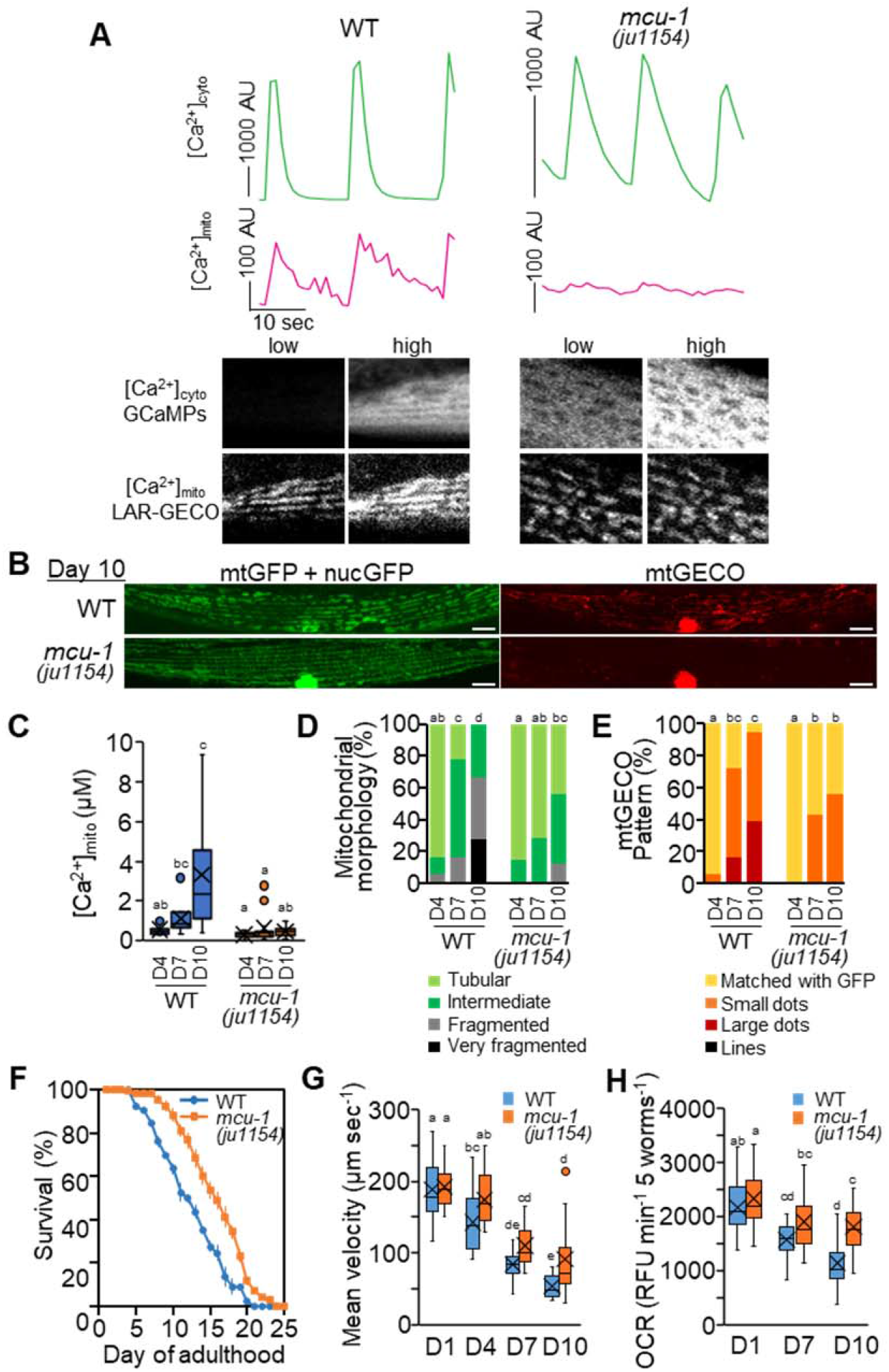
Effect of loss-of-function mutation in *mcu-1* on muscle aging in *C. elegans*. (A) Fluorescent signals of cytosolic GCaMPs ([Ca^2+^]_cyto_, green) and mtGECO ([Ca^2+^]_mito_, magenta) in body wall muscles of immobilized WT worm (*goeIs3; aceIs1*, left panel) or *mcu-1(ju1154)* mutant (right panel). Typical fluorescent images of muscle cytosolic GCaMP and mtGECO at low and high [Ca^2+^]_cyto_ levels were shown in the bottom panel. Mitochondrial Ca^2+^ uptake during cytosolic Ca^2+^ fluctuations was suppressed in *mcu-1* mutants. (B) Representative fluorescence images of mitochondria-targeted GFP (mtGFP), nuclear-targeted GFP (nucGFP), and mitochondrial Ca^2+^ probe (mtGECO) in body wall muscle cells in WT (*ccIs4251; aceIs1*) and *mcu-1(ju1154)* worms at D10 of adulthood. Mitochondrial fragmentation and mtGECO accumulation were abolished by *mcu-1* mutation. Fluorescence signals on the nuclei observed in the ‘mtGECO’ channel are from spectral bleed-through from nucGFP (See also Figure S1B). Scale bars, 10 μm. (C) Loss-of-function mutation in *mcu-1* decreased mitochondrial Ca^2+^ levels. Mitochondrial Ca^2+^ levels on mtGFP-positive mitochondria in muscle cells on D4, D7 and D10 of WT or *mcu-1(ju1154)* adulthood worms was calculated as described in Methods and Protocols (n=14). (D) Qualitative analysis of body wall muscle cells with abnormal mitochondrial morphology (n=18-20 muscle cells from at least 6 worms). (E) Qualitative analysis of body wall muscle cells with different mtGECO pattern (n=18-20 muscle cells from at least 6 worms). (F) Survival curve of WT and *mcu-1(ju1154)* worms. Data are presented as the mean ± standard error from four plates with 25-30 worms per plate. (G) Mean velocity of WT (*ccIs4251; aceIs1*) and *mcu-1(ju1154)* worms (n=16-22 worms). Loss-of-function mutation in *mcu-1* ameliorated age-related reductions in movement velocity. (H) The OCRs of WT and *mcu-1(ju1154)* mutant worms. OCR is expressed as the relative fluorescence unit (RFU) per minute per five worms (Materials and Protocols). Age-related reduction in OCR was improved in the mutants. Data represent the value from at least 10 repeats, each with 5 worms. Letters on the tops of bars indicate statistical significance by one-way ANOVA with Dunn’s multiple comparison test (C, G, H) or the chi-square test (D, E) (*p* < 0.05).

Intriguingly, the *mcu-1* mutant was shown to suppress mitochondrial fragmentation and severe loss of mitochondrial mass, as well as age-related Ca^2+^ accumulation in mitochondria (Figure 2B, C and D). As a result, the proportion of muscle cells displaying progressing mitophagy with age also decreased (Figure 2B and E). In the control wild-type (WT) with *aceIs1* and *ccIs4251* transgenes, approximately half of the population had died by 10-day-old adulthood (D10), with surviving individuals displaying significantly reduced motor activity (mean velocity) and oxygen consumption rate (OCR) activity compared to 1-day-old adulthood (D1) (Figures 2F, G and H). These age-related declines were significantly reduced in the *mcu-1(ju1154)* null mutation in the same background. In addition, in the muscle on D10 adults treated with *mcu-1* RNAi, less than 20% muscular cells were ‘fragmented’, [Ca^2+^]_mito_ did not significantly increase until 10-days old, and mitophagy activity in muscle cells was decreased (Supplementary Figure S3). The life span was also extended in the *mcu-1* mutant; the difference in life expectancy between the mutant and control was 3.2 days (control, 12.5 ± 0.2 days; *mcu-1(ju1154)*, 15.7 ± 0.4 days; *p* < 0.01) (Figure 2F). A similar result was observed between wild-type N2 and the mutant strain CZ19982 without the transgenic reporters; the original *mcu-1(ju1154)* mutation extended life span (N2, 14.7 ± 0.5 days; CZ19982 *mcu-1(ju1154)*, 17.6 ± 0.8 days; *p* < 0.05).

Recently, it has been reported that the highly conserved ryanodine receptor (RyR), UNC-68 in *C. elegans*, is oxidized with age which results in age-dependent ‘leaky’ channels^24^. Live imaging of cytosolic Ca^2+^ ([Ca^2+^]_cyto_) with contraction and relaxation of the body wall muscles was performed on D4, D10 and D15 of ATU2301. The result showed that the [Ca^2+^]_cyto_ peak width increased with age (Supplementary Figure S4). The full-width half-maximum (FWHM)^25^ was 5.8 ± 2.6 sec, 15.0 ± 16.9 sec, and 64.2 ± 35.9 sec in D4, D10, and D15 of adulthood, respectively. These observations suggest that age-related RyR dysfunction causing prolonged elevation of cytosolic Ca^2+^ may contribute to the increase in muscle [Ca^2+^]_mito_ with age.

### Effect of pharmacological inhibition of MCU-1with Ru360 on muscle aging in *C. elegans*

Having established that genetic ablations of *mcu-1* were sufficient to improve age-related muscle mitochondrial changes, we examined whether pharmacological inhibition of MCU-1 could similarly improve muscle health with age. Ru360, a specific mitochondrial calcium uptake inhibitor^26,27^, was used to inhibit mitochondrial Ca^2+^ influx. As this compound is significantly restricted in intact mammalian systems due to its poor cell permeability^26,27^, we first evaluated the penetration of Ru360 into intact *C. elegans* continuously cultured at a final concentration of 10 μM Ru360 from egg to adulthood. In adults of ATU2301 treated with Ru360, synchronized Ca^2+^ influx into the mitochondria was significantly lost (Figure 3A), as it was the *mcu-1* null mutant (Figure 2A), indicating that Ru360 is permeable to *C. elegans* muscle cells. As expected, Ru360 treatment prevented age-related changes in the body wall muscles of *C. elegans* including upregulated [Ca^2+^]_mito_, mitochondrial fragmentation and formation of mtGECO-structures (Figures 3B-E). Furthermore, Ru360 improved mobility with age (Figure 3F). These results confirm that inhibition of MCU-1 mediated Ca^2+^ influx attenuates age-dependent changes in mitochondrial morphology and Ca^2+^ levels, suppress the emergence of Ca^2+^-accumulated structures, and ultimately attenuates movement decline with age. The *mcu-1* mutant, *mcu-1* RNAi, and MCU-1 inhibitor studies combine in demonstrating that *C. elegans* muscle decline with age is normally driven by excessive Ca^2+^ influx into mitochondria and not merely elevated cytosolic Ca^2+^ levels due to leaky RyR channels^24^.

**Figure 3.**
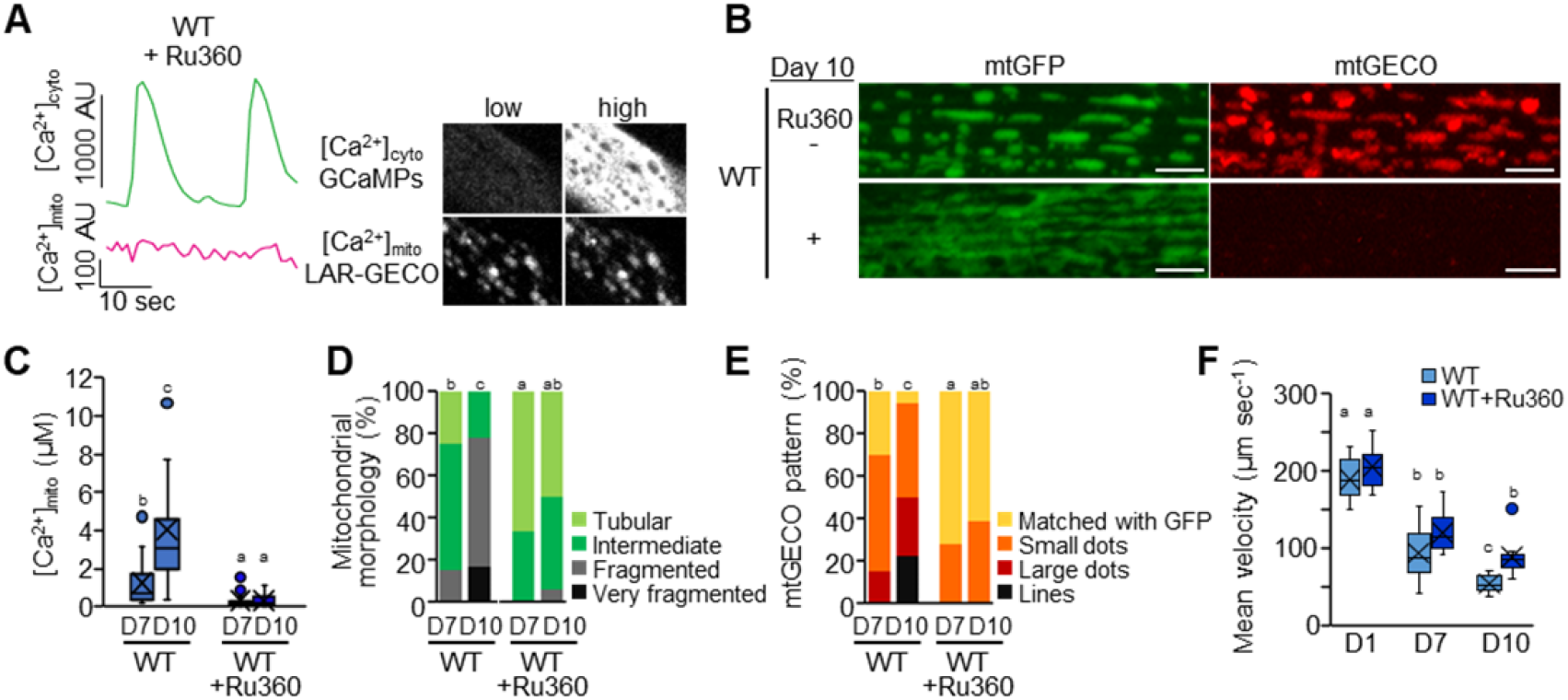
Pharmacological inhibition of *mcu-1* with Ru360 on muscle aging in *C. elegans*. (A) Suppression of mitochondrial Ca^2+^ uptake during cytosolic Ca^2+^ fluctuations by Ru360 treatment. Fluorescent signals of cytosolic GCaMPs ([Ca^2+^]_cyto_, green) and mtGECO ([Ca^2+^]_mito_, magenta) were monitored simultaneously in body wall muscles of Ru360-treated WT worm (*goeIs3; aceIs1*). Typical fluorescent images of muscle cytosolic GCaMP and mtGECO at low and high [Ca^2+^]_cyto_ levels were shown in the right panel. (B) Representative fluorescent images of mitochondria (mtGFP) and a mitochondrial Ca^2+^ probe (mtGECO) in body wall muscle cells in untreated and Ru360-treated WT worms (*ccIs4251; aceIs1*) at D10 of adulthood. Ru360 treatment suppressed age-related mitochondrial fragmentation and mtGECO accumulation. Scale bars, 5 μm. (C) Mitochondrial Ca^2+^ levels in mtGFP-positive mitochondria of muscle cells on D7 and D10 of adulthood. The levels were calculated as described in Methods and Protocols (n=14). Ru360 treatment restrained the age-related increase in mitochondrial Ca^2+^ levels. (D) Qualitative analysis of body wall muscle cells with abnormal mitochondrial morphology (n=18 muscle cells from at least 6 worms). (E) Qualitative analysis of body wall muscle cells with different mtGECO pattern (n=18 muscle cells from at least 6 worms). (F) Mean velocity of untreated and Ru360-treated WT worms (*ccIs4251; aceIs1*) (n=10-13 worms). Ru360 treatment improved the age-related decline in mean velocity. Letters on the tops of bars indicate statistical significance by one-way ANOVA with Dunn’s multiple comparison test (C, F) or the chi-square test (D, E) (*p* < 0.05).

### Improvement of health by inhibition of *mcu-1* in the *C. elegans* DMD model

In the *C. elegans* DMD model *(dys-1(eg33)*, altered calcium homeostasis in muscle is a primary pathology. These dystrophin mutants display increased mitochondrial fragmentation in muscle cells, and decreased mobility^17,18^. Given the positive effects of inhibition of *mcu-1* on muscle calcium homeostasis and health with age, we were curious if Ru360 could have beneficial effects on the dystrophy model. After immobilization with polystyrene microspheres, cytosolic Ca^2+^ live imaging was performed on single muscle cells of D4 animals (Figure 4A). The FWHM duration of the *dys-1*(*eg33*) mutant, was significantly (23.0 ± 13.5 sec) longer than that of the wild type (8.0 ± 0.8 sec), suggesting that muscle rigidity occurred (Figure 4A and B). FWHM broadened in *dys-1(eg33)* D4 animals and was similar to WT aged animals. In addition, [Ca^2+^]_mito_ also maintained higher levels in mutant muscle cells (Figure 4A). Intriguingly, Ru360 treatment not only suppressed Ca^2+^ influx into mitochondria, but also significantly improved the broadening of FWHM (11.6 ± 7.7 sec) in the *dys-1* mutants (Figure 4 A and B). We also found that fragmentation of mitochondria and Ca^2+^ accumulation in mitochondria in the muscle cells of dystrophin mutant worms were decreased by Ru360 treatment (Figures 4C and D).

**Figure 4.**
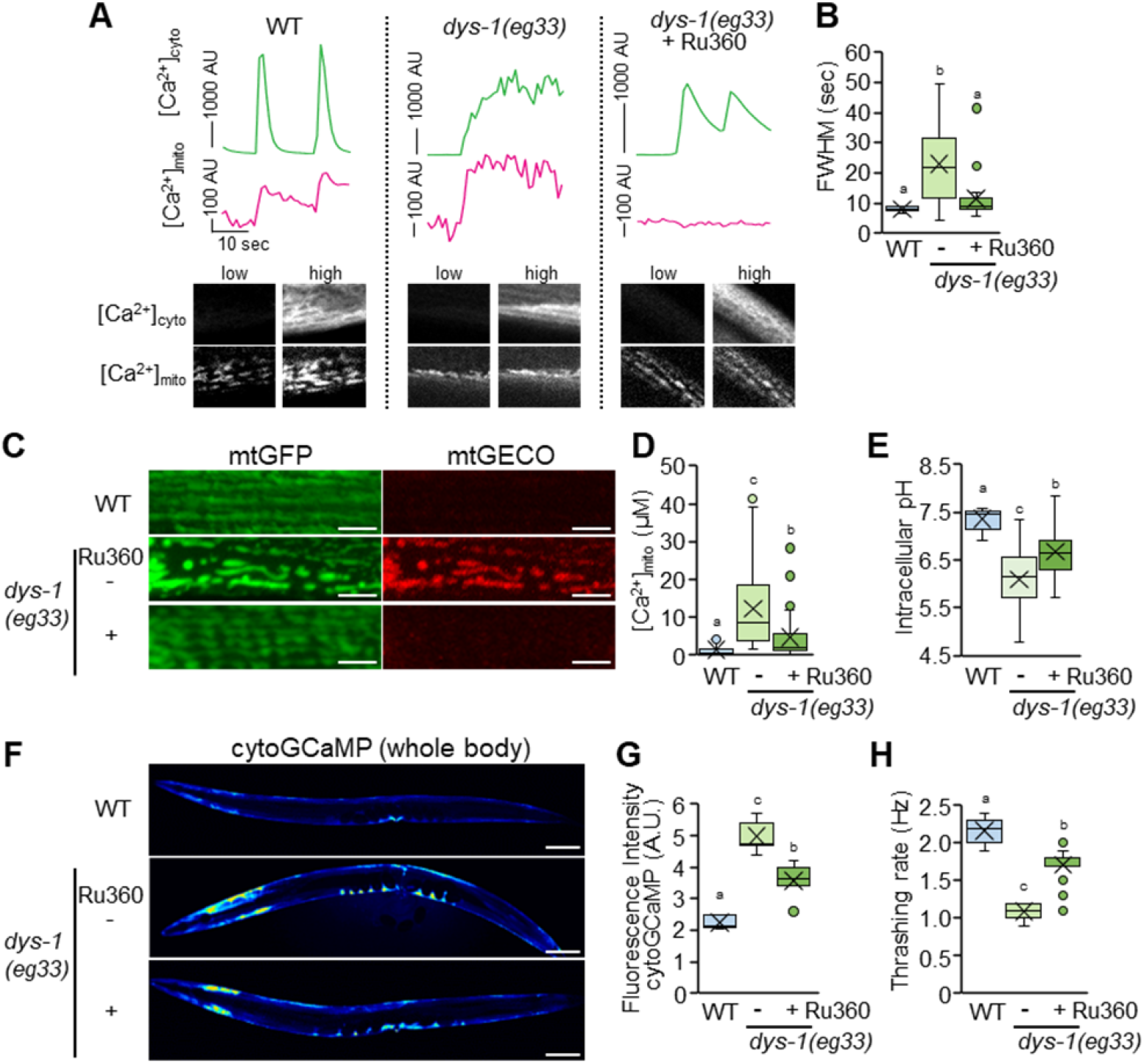
Improvement of dystrophy model by inhibition of *mcu-1* in *C. elegans*. (A) Mitochondrial Ca^2+^ uptake (magenta) during spontaneous cytosolic Ca^2+^ fluctuations (green) in body wall muscles of immobilized WT worm (*goeIs3; aceIs1*, left panel), *dys-1(eg33)* (center panel), and Ru360-treated *dys-1(eg33)* (right panel). Fluorescent signals of cytosolic GCaMPs and mtGECO were monitored simultaneously. Typical fluorescent images of muscle cytosolic GCaMP and mtGECO at low and high levels were shown in the bottom panel. (B) Full-width half-maximum (FWHM) of the [Ca^2+^]_cyto_ peaks in WT (*goeIs3; aceIs1*), untreated and Ru360-treated *dys-1(eg33)* mutant worms (n=16-20). (C) Representative fluorescent images of mitochondria and mtGECO in the body wall muscle cells. Scale bars, 5 μm. (D) Mitochondrial Ca^2+^ levels in mtGFP-positive mitochondria of muscle cells on D2 of adulthood. The levels were calculated as described in Methods and Protocols (n=31-39). (E) Intracellular pH in body wall muscle cells in WT (N2) and untreated and Ru360-treated *dys-1(eg33)* mutant worms on D3 of adulthood (n=11-13). (F and G) Fluorescent images (F) and fluorescence intensity (G) of the whole body of transgenic *C. elegans* expressing a cytosolic Ca^2+^ probe (GCaMP) in body wall muscle cells. The *dys-1(eg33)* mutant exhibited higher fluorescence intensity for cytosolic GCaMP than the WT worms, and Ru360 treatment rescued this change. Scale bars, 100 μm. (H) Thrashing rate (Hz) in WT (*goeIs3; aceIs1*), untreated and Ru360-treated *dys-1(eg33)* mutant worms. Inhibition of mitochondrial Ca^2+^ uptake by Ru360 improved the decline in motor activity (thrashing) of the *dys-1(eg33)* mutant (n=25). Letters on the tops of bars indicate statistical significance by one-way ANOVA with Dunn’s multiple comparison test (B, D, E, G, H) (*p* < 0.05).

In both mammalian cells and *C. elegans* body wall muscle cells, intracellular acidification is caused by mitochondrial fragmentation and the pH drops from 7.5 to about 7.0^28^. Although measured with different pH indicators, we obtained similar results with wild-type pH 7.4 versus pH 6.1 in the *dys-1* mutant and pH 6.7 in Ru360 treated *dys-1* mutants (Figure 4E). In addition, we monitored Ca^2+^ level in the cytosol of the muscle cells on D2 of adulthood after anesthesia with sodium azide. Compared with the WT counterparts, the dystrophin mutants had significantly higher cytosolic Ca^2+^ levels, and surprisingly, Ru360 treatment decreased cytosolic Ca^2+^ level in dystrophin mutants (Figure 4F and G). Furthermore, Ru360 improved mobility in the *dys-1* mutants (Figure 4H). Similarly, to Ru360 treatment, *mcu-1* RNAi prevented the progression of mitochondrial Ca^2+^ accumulation and decline in movement of dystrophin mutant worms (Supplementary Figure S5). These results indicate that suppression of mitochondrial Ca^2+^ influx improves muscular Ca^2+^ homeostasis in the *dys-1*(*eg33*) mutant and restores the motility dysfunction. Thus, elevated mitochondrial Ca^2+^ causes impaired muscle health not only with age but also in DMD.

In MCU knockout mice, skeletal muscle showed altered phosphorylation and activity of pyruvate dehydrogenase, significantly impairing the ability to perform strenuous work^29^. Rizzuto’s group also shows that MCU silencing causes muscle atrophy^10^. In contrast, our present study shows that MCU inhibition (genetic ablation of *mcu-1* and Ru360 treatment) ameliorated muscular function decline with age and DMD in *C. elegans*. Recently, MCU-1 inhibitors, which are also ruthenium compounds with improved *in vivo* permeability such as Ruthenium Red and Ru265, have been developed and are being investigated as potential therapeutic agents for cardiac dysfunctions^30,31^. These findings suggest MCU has an evolutionarily conserved role in muscle health. Further, differences in the role of MCU function with development and growth vs. aging and pathology are now apparent. Likewise, mutations in an MCU regulator MICU1, which increases resting mitochondrial Ca^2+^ levels, caused neuromuscular disorders with cognitive decline, muscle weakness, and an extrapyramidal motor disorder^11^. In particular, the aging state in mammalian muscles, where muscle satellite cells are gradually lost and the regenerative capacity is reduced^32^, is highly similar to the aging of body wall muscle cells in adult *C. elegans*. All together suggest that controlling MCU function can be the potential target for diagnosis of sarcopenia even in the mammalian system.

## Conclusion

Here we show that the blockage of MCU-1function by genetic or pharmacological modulation improves health in both aging *C. elegans* and in *C. elegans* with muscular dystrophy. These observations suggest that loss of calcium homeostasis is an early event in muscle aging that can be mitigated against by improving mitochondrial calcium buffering capacity. Suppression of mitochondrial Ca^2+^ influx prevented the formation of Ca^2+^-accumulated structures in body wall muscle cells. These age associated Ca^2+^-accumulations appear to normally be removed via mitophagy. These results suggest that RyR, which causes increased Ca^2+^ with age, MCU-1, which facilitates increase mitochondrial Ca^2+^ with age, and mitophagy, which maintains mitochondrial homeostasis, are part of a coordinated system that fails to maintain muscle health with age and which may be targeted for improved muscle health.

## Acknowledgements

We acknowledge the CGC, funded by the U.S. National Institutes of Health (NIH) Office of Research Infrastructure Program (P40 OD010440), for providing the *C. elegans* strains. We acknowledge Dr. Julie Ahringer and Source BioScience for providing the RNAi library. L4440 was a gift from Andrew Fire (Addgene plasmid # 1654 ; http://n2t.net/addgene:1654;RRID:Addgene_1654). This work was funded in part by Advanced Research and Development Programs for Medical Innovation, AMED-CREST (16814305), JSPS KAKENHI grant numbers JP15H05937, JP19H04054 and JP21K06149, BBSRC (Grants BB/N015894/1 and BB/P025781/1), and the UK Space Agency and the STFC (Grant ST/S006338/1). This work was supported by AMED Grant Number JP21zf0127001.

## Author contributions

A.H. and T.K. conceived and designed the study. M.T., Y.N., Y.I., S.S., T.K. and A.H. conducted experiments and analysed the data. Y.K. contributed to the generation of transgenic *C. elegans*. A.H., M.T., S.S., N.J.S. and T.K. wrote the manuscript. N.J.S. and T.A. supervised the mitochondrial dysfunction project and chemical treatment. All authors read and approved the final paper.

## Materials and Methods

### Reagents and Tools table

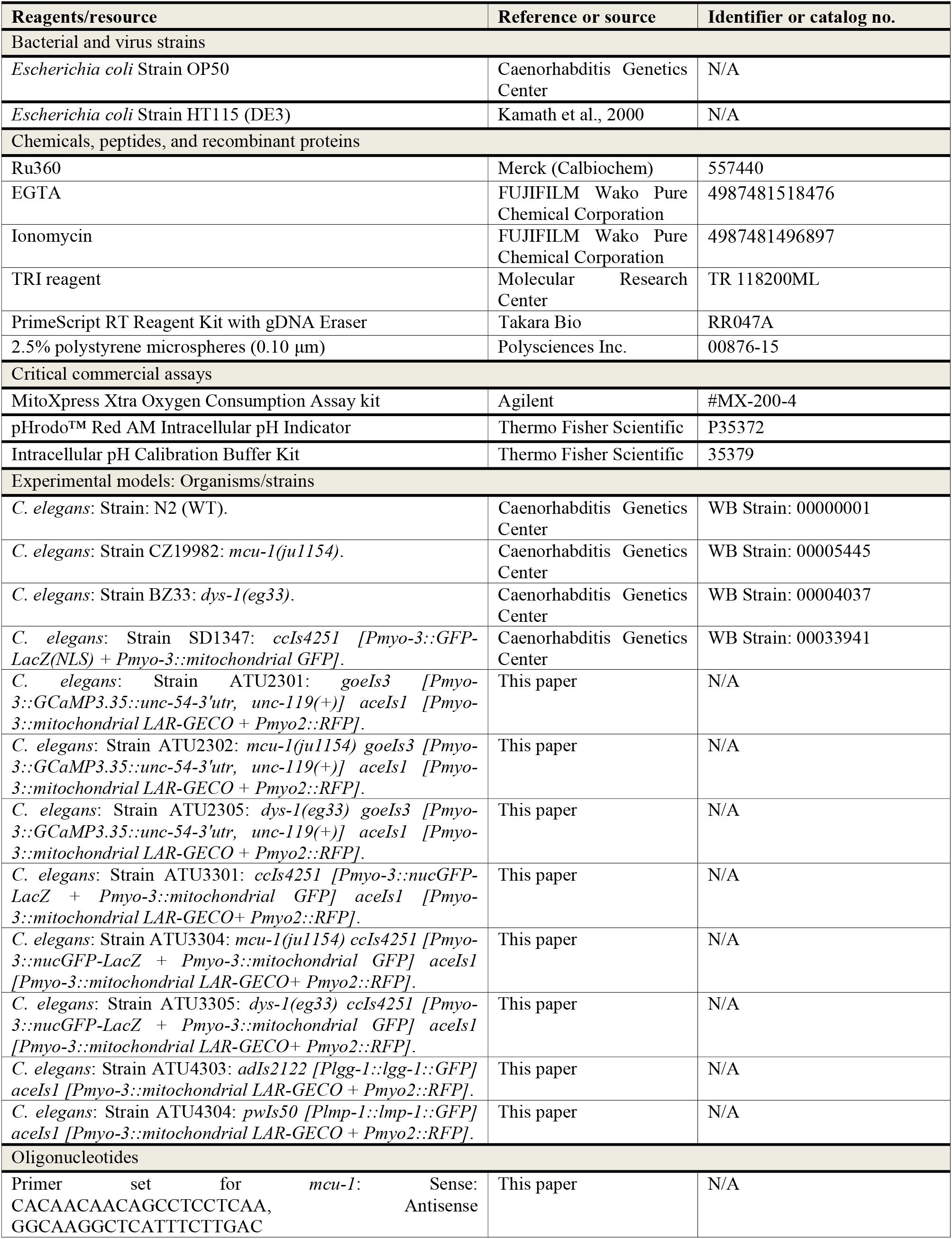
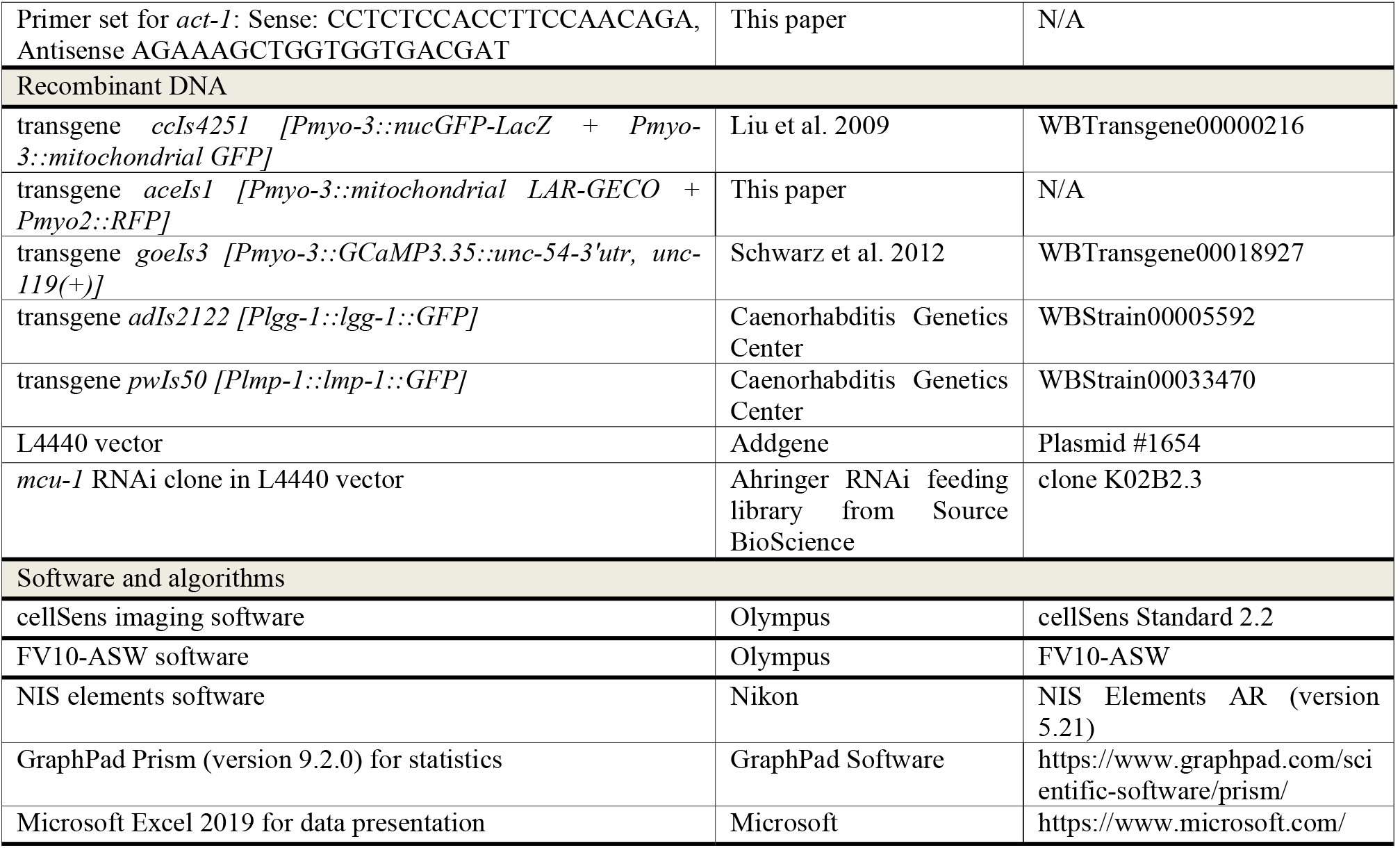

## Methods and Protocols

### Culture conditions

We followed standard procedures for *C. elegans* strain maintenance^33^. All strains were cultured on nematode growth medium (NGM) plates with OP50 as a food source at 20°C. The worms were synchronized by egg laying for 3h.

### Imaging of mitochondrial morphology, mitochondrial Ca^2+^, cytosolic Ca^2+^ and nuclear morphology

Mitochondrial morphology and mitochondrial Ca^2+^ in body wall muscle cells were observed by assaying expression of the transgenes *ccIs4251 [Pmyo-3::nucGFP-LacZ + Pmyo-3::mitochondrial GFP]*^21^, *goeIs3 [myo-3p::SL1::GCamP3*.*35::SL2::unc54 3’UTR + unc-119(+)]*^22^ and *aceIs1 [Pmyo-3::mitochondrial LAR-GECO + Pmyo2::RFP]*, respectively. Synchronized worms were mounted on a microscope slide with a 6.5-mm square, 20-μm deep well made with a water-repellent coating (Matsunami Glass Ind., Ltd.) with a 100 mM NaN_3_ solution (Z-stack imaging) or 2.5% polystyrene microspheres (0.10 μm, Polysciences Inc.) (time-lapse confocal imaging). Z-stack images and time-lapse confocal images of GFP and mtGECO fluorescence were observed by an FV10i confocal laser-scanning microscope (Olympus). Time-lapse confocal images of cytosolic GCaMP and mtGECO fluorescence in body wall muscle cells were acquired at room temperature (20 ∼ 22 °C) on a CSU-W1 spinning disk scanner (single camera split-view model, Yokogawa Electric Corporation) on an Eclipse Ti2-E inverted microscope (Nikon) with a CFI Apo TIRF 60x N.A. 1.49 objective (Nikon). Worms were simultaneously illuminated by two laser lines at 488 nm (Sapphire 488, Coherent) and 561 nm (OBIS, Coherent). Emission fluorescence of GCaMP and mtGECO was divided by a dichroic mirror (561LP, Semrock, Inc.) and projected onto adjacent halves of an EMCCD camera (iXon Life 888; Andor Technology). Images were acquired every 1 sec for calcium imaging and analyzed using of NIS-Elements AR software (Nikon).

Mitochondria were grouped into categories by morphology according to Regmi et al^14^ as follows: ‘tubular’; images indicating a majority of long interconnected mitochondrial networks, ‘intermediate’; images indicating a combination of interconnected mitochondrial networks along with some smaller fragmented mitochondria, ‘fragmented’; images indicating a majority of short mitochondria, ‘very fragmented’; images indicating sparse small round mitochondria. The morphological categories of mtGECO were grouped into categories as follows: ‘matched with GFP’; images that match with mitochondrial morphology were classified as matched with GFP, ‘small dots’; images indicating more than 10 dots per cell smaller than 1 μm were classified as small dots, ‘large dots’; images indicating more than 10 dots per cell larger than 1 μm were classified as large dots, ‘lines’; images indicating more than 5 linear signals per cell were classified as lines.

### Measurement of mitochondrial Ca^2+^ and cytosolic Ca^2+^ levels

Mitochondrial Ca^2+^ and cytosolic Ca^2+^ in body wall muscle cells were observed by assaying expression of the transgenes *aceIs1 [Pmyo-3::mitochondrial LAR-GECO + Pmyo2::RFP]* and *goeIs3 [Pmyo-3::GCaMP3*.*35::unc-54-3’utr, unc-119(+)]*^22^, respectively. The fluorescent signals of mtGECO in a constant area and cytosolic GCaMP in the whole body were imaged by an FV10i confocal laser-scanning microscope (Olympus). The Ca^2+^ concentration in muscle mitochondria ([Ca^2+^]_mito_) was calculated using the following equation^34^.

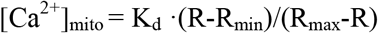

Where K_d_ (12 μM) indicates the dissociation constant between Ca^2+^ and the LAR-GECO probe^20^ and R indicates the ratio of fluorescence intensity of mtGECO to that of mtGFP in a constant area. Muscle mitochondria at each age were exteriorized in the incision of worms cut by a blade and calibrated in the presence of 10 mM EGTA (R_min_) or by the addition with 5 μM Ionomycin in 1 mM CaCl_2_ (R_max_) (Supplementary Figure S2A and B).

### Measurement of life span

To measure life span, the living and dead worms were counted every day. A total of 100-120 worms were placed on four replicate plates, with 25-30 worms per plate. The worms were transferred to a fresh plate every 2 to 3 days.

### Measurement of mean velocity and locomotion activity

Synchronized worms were transferred to NGM plates with no bacteria. Movement was recorded by using stereomicroscopy (SMZ18; Nikon), a device camera (DP74; Olympus), and imaging software (cellSens Standard 2.2; Olympus). The distance moved every five seconds was measured, and the average of four points was calculated. To determine locomotion activity, we calculated the average thrashing rate for after tapping in liquid culture for 10 s^17,35^.

### Measurement of the OCR

Synchronized worms were washed three times with M9 buffer, and 5 worms per well were transferred to a 96-well, black wall, clear bottom plate containing 45 μl of M9 buffer. Five microliters of reagent from a MitoXpress Xtra Oxygen Consumption Assay kit (Agilent) was added per well and mixed. The wells were covered with mineral Oil, and fluorescence was detected with a microplate reader (Spark 10M; Tecan). Each experiment was performed in at least 10 wells using different worms.

### RNAi treatment

For *mcu-1* RNAi, clones from the Ahringer RNAi feeding library (Source BioScience, Nottingham, UK) were used. The clone number was K02B2.3. RNAi was performed by bacterial feeding as described by Kamath et al^36^. L4 larvae from the WT (*ccIs4251*; *aceIs1*) and *dys-1(eg33)* were transferred to NGM plates coated with HT115(DE3) bacteria expressing dsRNA for *mcu-1* gene. Bacteria containing the empty L4440 vector were used as a control. The worms were transferred to newly prepared RNAi plates every 2 days.

### Real-time quantitative RT-PCR

Total RNA was extracted from 25-30 worms on D7 of adulthood with TRI reagent (Molecular Research Center). cDNA was synthesized using a PrimeScript RT Reagent Kit with gDNA Eraser (Takara Bio Inc.). Data was normalized against the expression of *act-1*. The experiment was repeated 3 times from different plates.

### Ru360 treatment

Ru360 (Merck) was dissolved in water and used at a final concentration of 10 μM. Eggs were laid on medium containing Ru360 and grown.

### Measurement of pH in body wall muscle cells

Intracellular pH of body wall muscles cells was measured by Invitrogen™ pHrodo™ Red AM Intracellular pH indicator (P35372, Thermo Fisher Scientific). Wild type N2 and untreated and Ru360-treated *dys-1(eg33)* mutant worms were incubated with 5 µM pHrodo™ Red AM Intracellular pH Indicator for 30 minutes at room temperature. Images were detected by an FV10i confocal laser-scanning microscope. The pH was determined by a standard curve using Intracellular pH Calibration Buffer Kit (P35379, Thermo Fisher Scientific).

### Statistical analysis

GraphPad Prism 9 software was used to determine statistical significance (GraphPad Software). Statistical analyses were performed using Student *t*-test, one-way ANOVA with Dunn’s multiple comparison test or the chi-square test.

### Strains, data and code availability

All experimental data of this study are available from the authors upon reasonable request. The ATU2301 and ATU3301 is available from the Caenorhabditis Genetic Center (CGC). The other ATU series of nematode strains constructed in this study is available from the authors in accordance with the Material Transfer Agreement. No new custom computer code or algorithm was used to generate the results reported in the paper. Any additional information required to reanalyze the data reported in this paper is available upon request.

## Supplemental information

**Figure S1.**
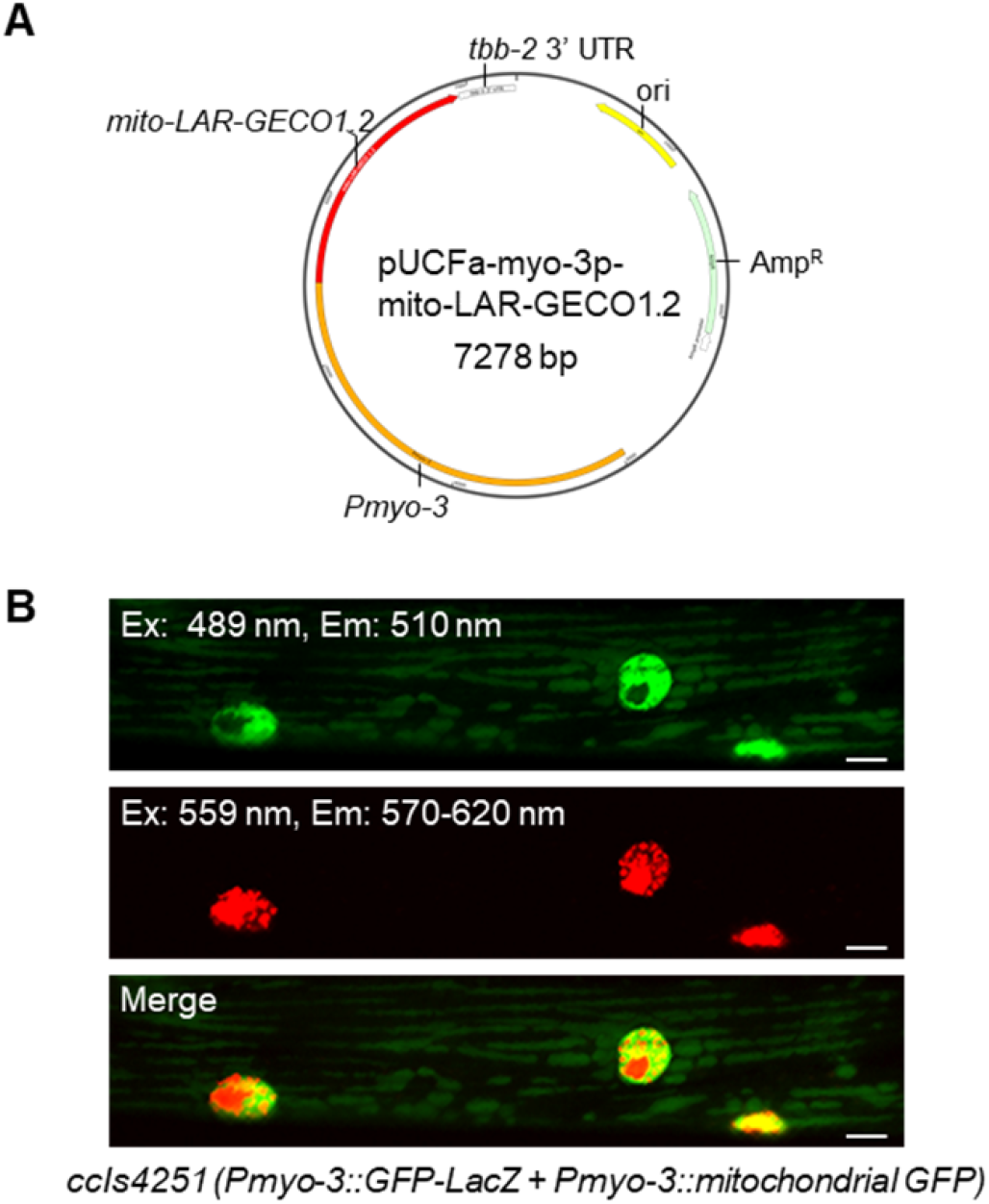
The construct of mitochondrial-targeted LAR-GECO1.2 in *aceIs1* and fluorescence of *ccIs4251* in body wall muscle cells. (A) Plasmid map of the construct used to introduce the *aceIs1* transgene. Mitochondrial-targeted LAR-GECO1.2 was constructed under the *myo-3* promoter with the *tbb-2* 3’ UTR. The LAR-GECO 1.2 codon was optimized for expression in *C. elegans*. pUC19 was used as a vector backbone. (B) The *ccIs4251* transgene showed not only GFP but also red fluorescence (not merged with GFP) in the muscle nuclei. *C. elegans* (*ccIs4251* [*Pmyo-3*::*GFP-LacZ(NLS)* + *Pmyo-3*::*mitochondrial GFP*]) whose body wall muscle cells expressed mitochondria-targeted GFP and nuclear-targeted GFP were monitored. The *ccIs4251* transgene shows red fluorescence in the muscle nuclei under the detection conditions for mtGECO (excitation: 559 nm, emission: 570-620 nm: FV10i confocal laser-scanning microscope, Olympus). Scale bars, 5 μm.

**Figure S2.**
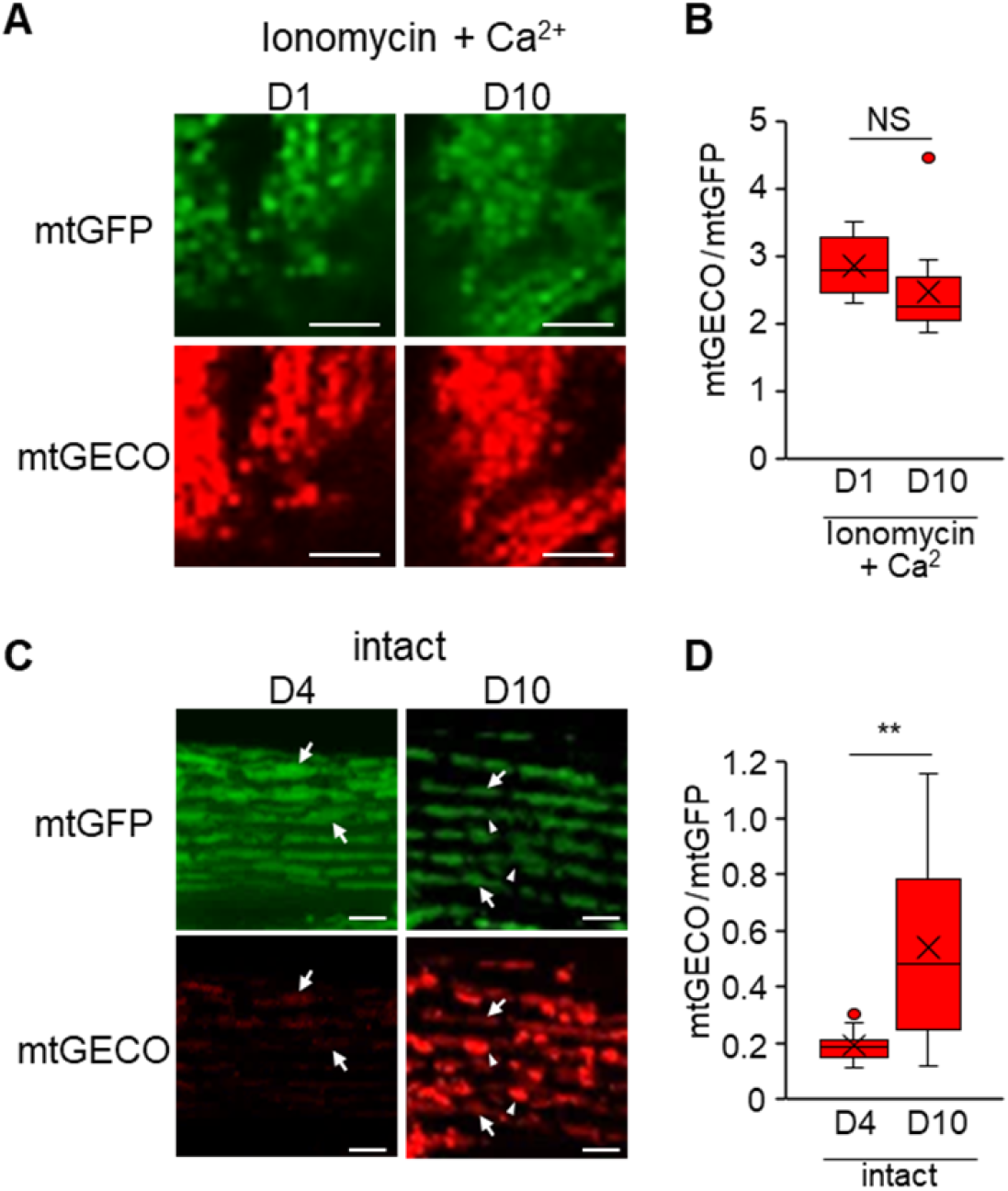
Measurement of mitochondrial Ca^2+^ levels in body wall muscle cells. (A) Representative fluorescence images of mtGFP and mtGECO in the mitochondria of body wall muscle cells in WT worms (*ccIs4251; aceIs1*). Muscle mitochondria at D1 and D10 of adulthood were exteriorized in the incision of worms cut by a blade and mixed with Ionomycin in CaCl_2_. Scale bars, 5 μm. (B) The ratio of fluorescence intensity of mtGECO to mtGFP with Ionomycin in CaCl_2_. No significant difference was observed between D1 and D10. NS means not significant (*p* > 0.05, Student’s *t*-test) (n=6-13). (C) Representative fluorescence images of mtGFP and mtGECO in the intact body wall muscle cells in WT worms (*ccIs4251; aceIs1*). The arrows indicate the typical position where the ratio of mtGECO to mtGFP was detected. The arrowheads indicate the position of mtGECO structures that no longer colocalized with mtGFP. Scale bars, 5 μm. (D) The ratio of fluorescence intensity of mtGECO to mtGFP in the intact body wall muscle cells. The value was significantly different between D4 and D10 (** *p* < 0.01, *t*-test) (n=19-21).

**Figure S3.**
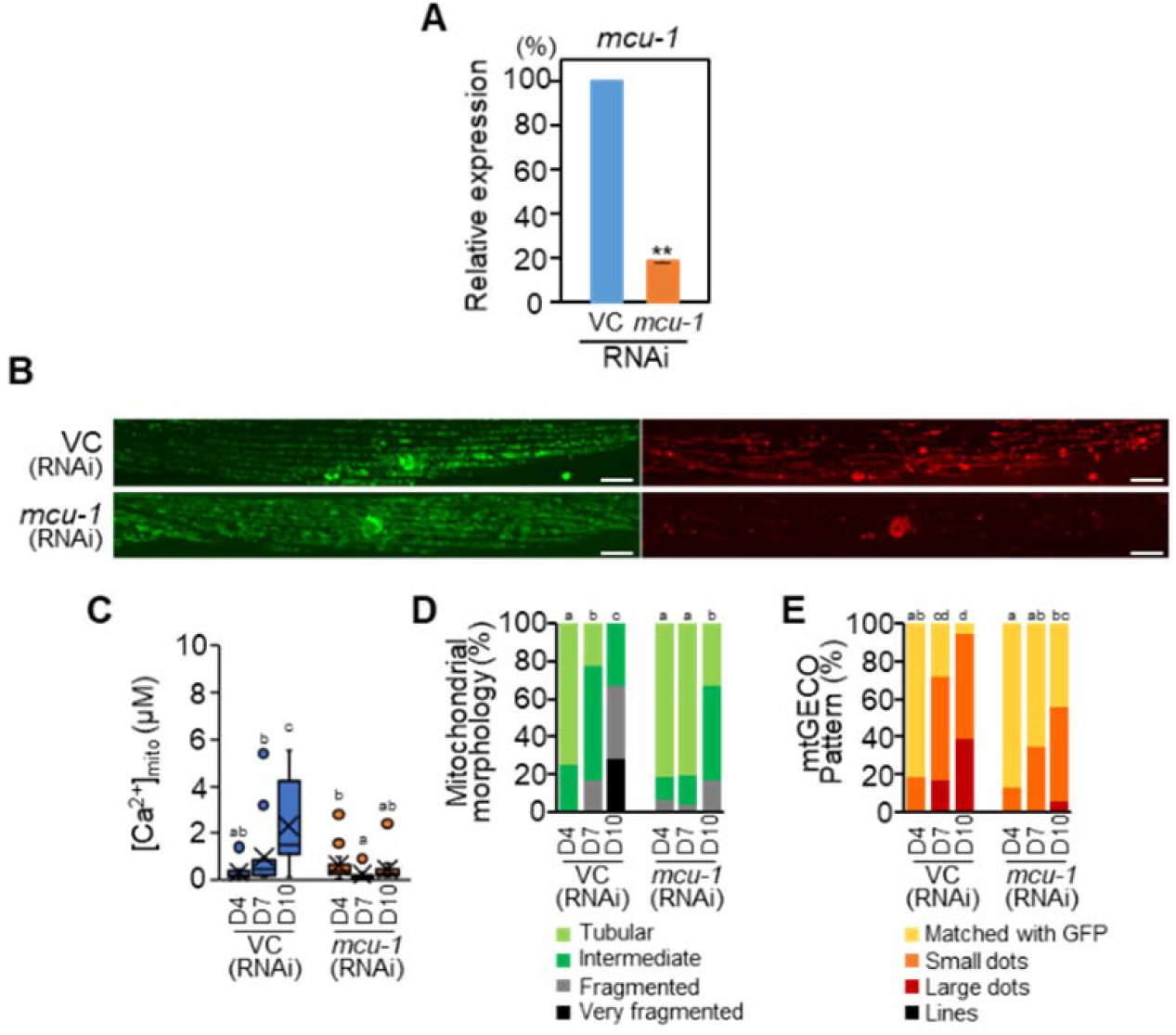
Effect of *mcu-1* RNAi on muscle aging in *C. elegans*. (A) The efficiency of RNAi treatment. WT worms (*ccIs4251; aceIs1*) were treated with a RNAi sequence (VC: vector control or *mcu-1*) from L4 larvae. Total RNA was extracted on D7 of adulthood, and RT-PCR was performed using sequence-specific primers. The data indicate a relative value with the value of VC set to 100%. Error bars represent standard errors among the four plates with 25-30 worms per plate (** *p* < 0.01, Student’s *t*-test). (B) Representative fluorescence images of mitochondria-targeted GFP (mtGFP), nuclear-targeted GFP (nucGFP), and mitochondrial Ca^2+^ probe (mtGECO) in body wall muscle cells of RNAi treated WT worms (*ccIs4251; aceIs1*) (VC and *mcu-1*) on D10 of adulthood. Scale bars, 10 μm. (C) Mitochondrial Ca^2+^ levels on mtGFP-positive mitochondria in muscle cells on D4, D7 and D10 adulthood worms treated with RNAi sequence was calculated as described in Methods and Protocols (n=14). (D) Qualitative analysis of body wall muscle cells with abnormal mitochondrial morphology (n=18-20 muscle cells from at least 6 worms). (E) Qualitative analysis of body wall muscle cells with different mtGECO pattern (n=18-20 muscle cells from at least 6 worms). Letters on the tops of bars indicate statistical significance by one-way ANOVA with Dunn’s multiple comparison test (C) or the chi-square test (D, E) (*p* < 0.05).

**Figure S4.**
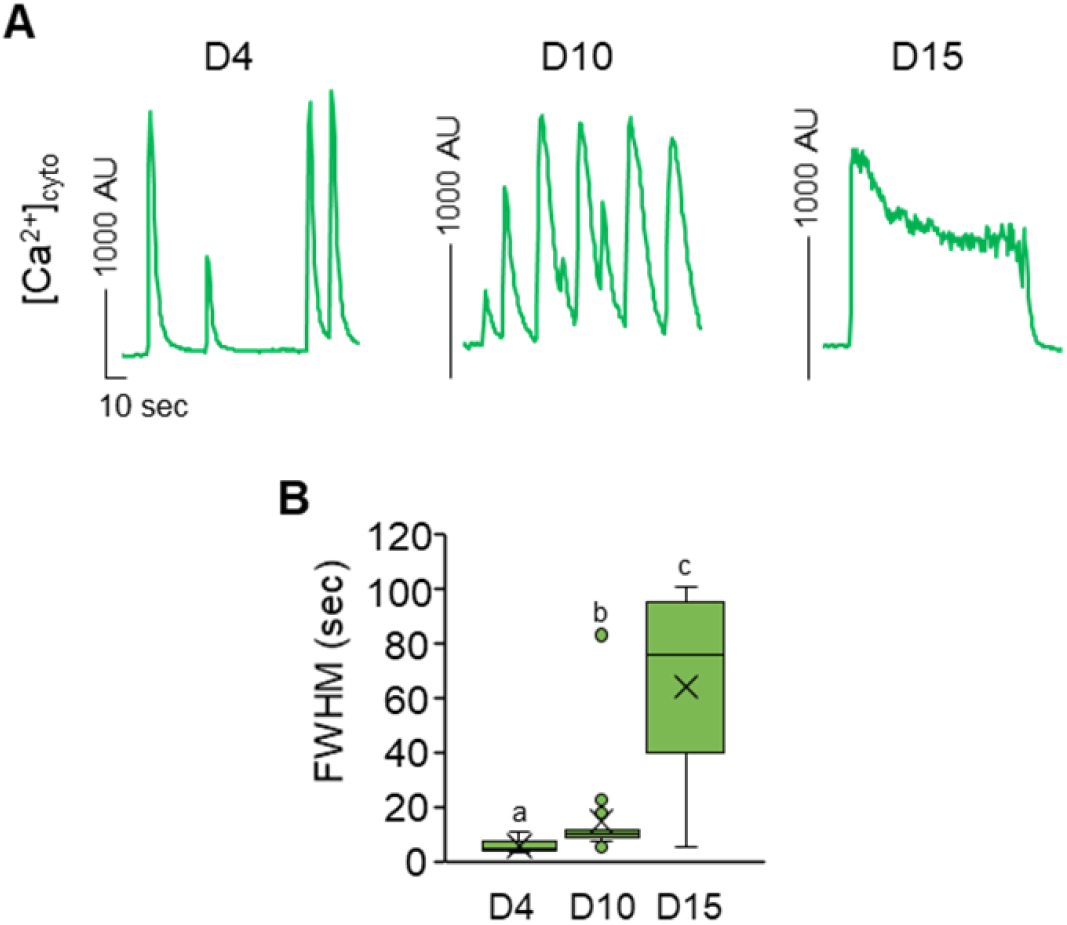
Age-related increase in cytosolic Ca^2+^ level in body wall muscle cells. (A) Fluorescent signals of cytosolic GCaMPs ([Ca^2+^]_cyto_) in body wall muscles of immobilized WT worm (*goeIs3; aceIs1*) at D4, D10 and D15 of adulthood. The peak width increased with age. (B) Full-width half-maximum (FWHM) of the [Ca^2+^]_cyto_ peaks in WT at D4, D10 and D15 of adulthood (n=10-18). FWHM increased with age. Letters on the tops of bars indicate statistical significance by one-way ANOVA with Dunn’s multiple comparison test (*p* < 0.05).

**Figure S5.**
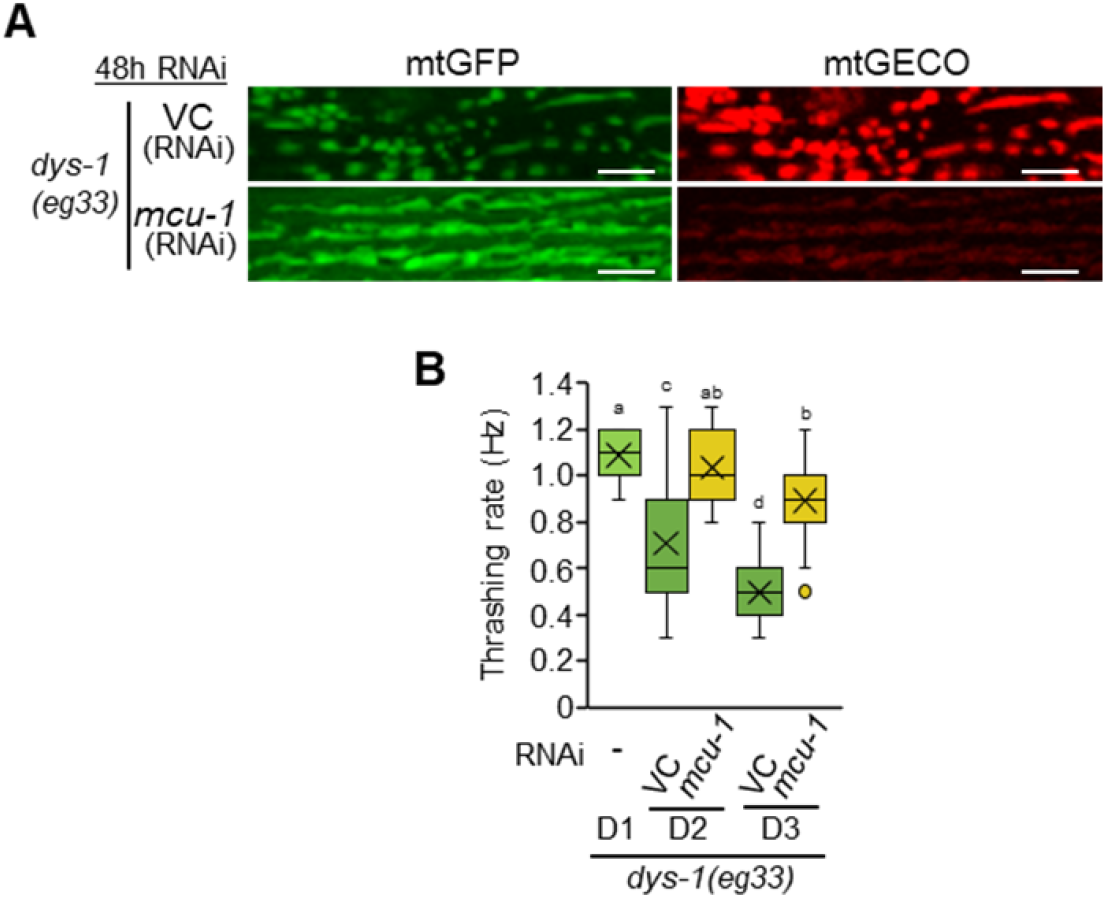
Improvement of dystrophy model by *mcu-1* RNAi in *C. elegans*. (A) Representative fluorescence images of mtGFP and mtGECO in the body wall muscle cells in the *dys-1(eg33)* mutant after 48 h of RNAi treatment (VC: vector control, and *mcu-1*). Mitochondrial fragmentation and mtGECO accumulation were abolished by *mcu-1* knockdown. Scale bars, 5 μm. (B) Thrashing rate (Hz) in RNAi treated *dys-1(eg33)* mutant worms. Inhibition of mitochondrial Ca^2+^ uptake by *mcu-1* RNAi improved the decline in motor activity (thrashing) of the *dys-1(eg33)* mutant (n=25). Letters on the tops of bars indicate statistical significance by one-way ANOVA with Dunn’s multiple comparison test (*p* < 0.05).

**Supporting Movie 1.**
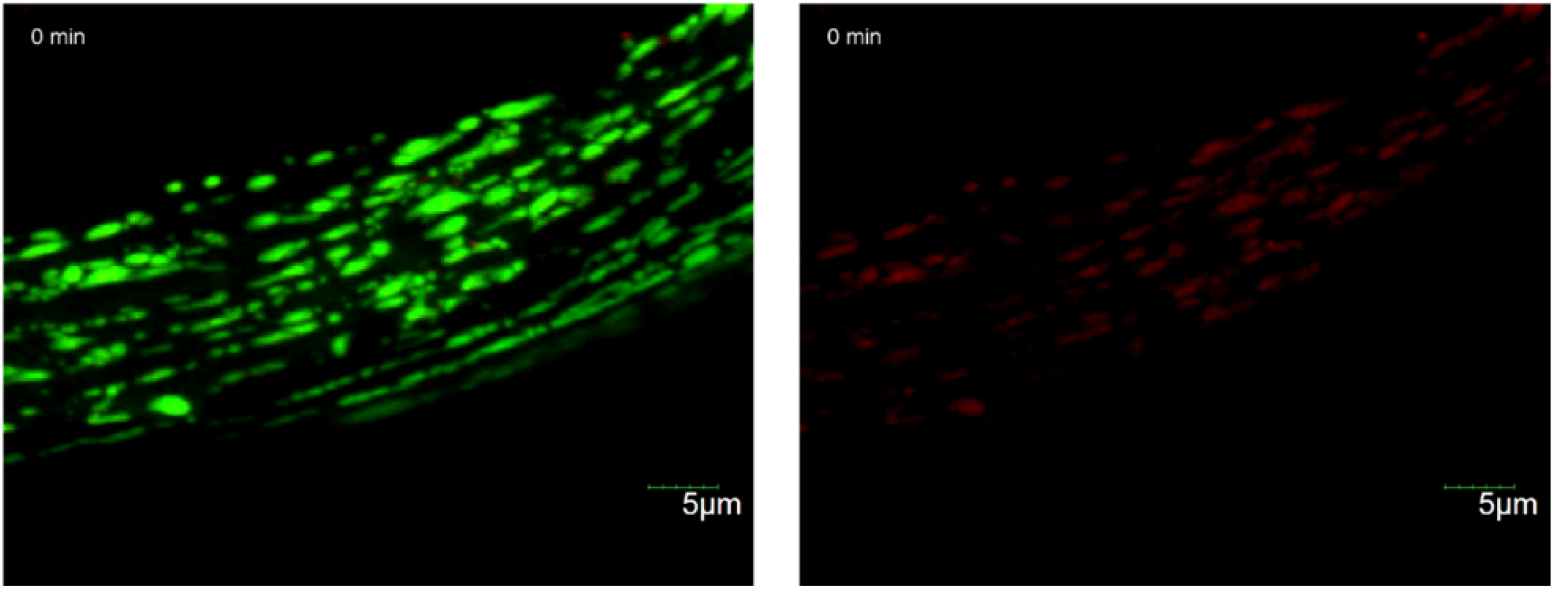
mtGECO structures were excised from mitochondria in D10 adult muscles. Time-lapse confocal fluorescent images of mitochondrial GFP (mtGFP, green) and mito-LAR-GECO1.2 (mtGECO, red) in body wall muscles of immobilized D10 WT worm. Images were acquired every 1 min for 20 min. Playback, 1 fps. Relates to Fig. 1.

**Supporting Movie 2.**
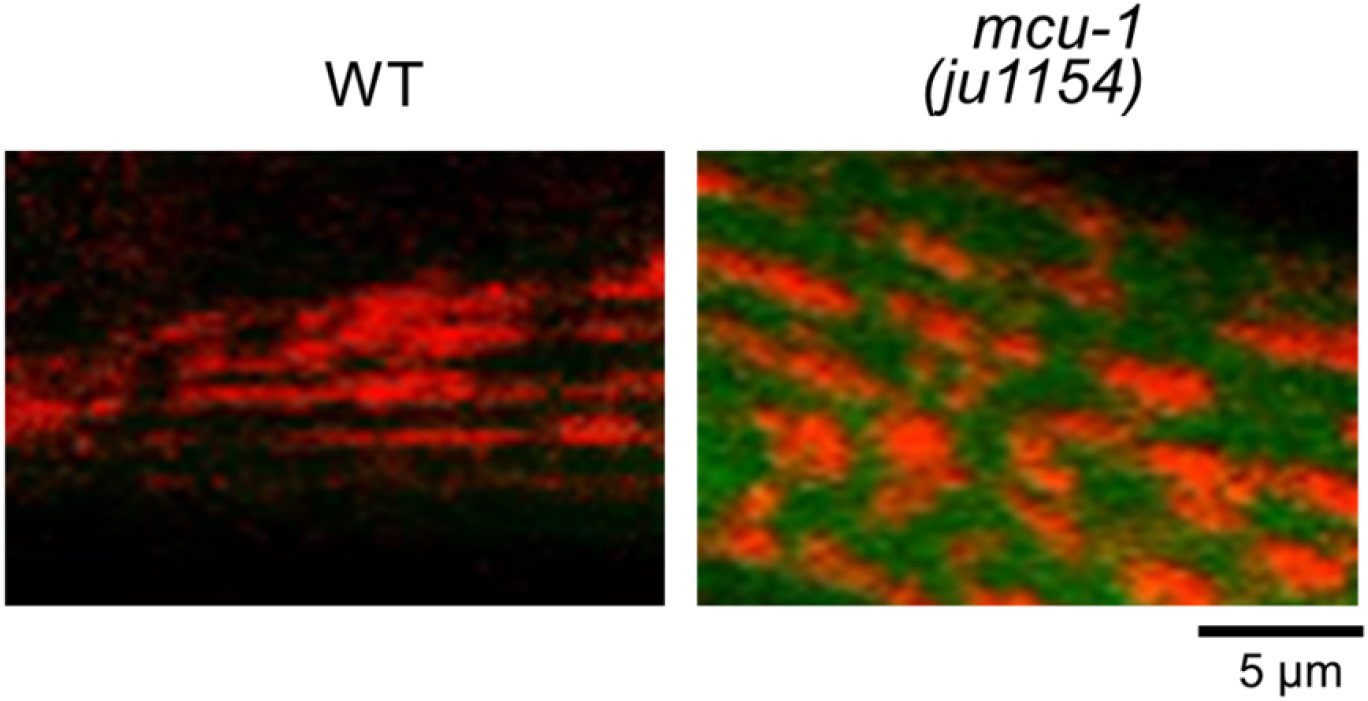
Time-lapse confocal fluorescent images of cytosolic GCaMPs ([Ca^2+^]_cyto_, green) and mtGECO ([Ca^2+^]_mito_, red) in body wall muscles of immobilized WT worm (*goeIs3; aceIs1*, left panel) or *mcu-1(ju1154)* mutant (right panel), showing that mitochondrial Ca^2+^ uptake during cytosolic Ca^2+^ fluctuations was blocked in *mcu-1* mutants. Images were acquired every 1 sec for 1 min. Playback, 10 fps. Relates to Fig. 2A.

**Supporting Movie 3.**
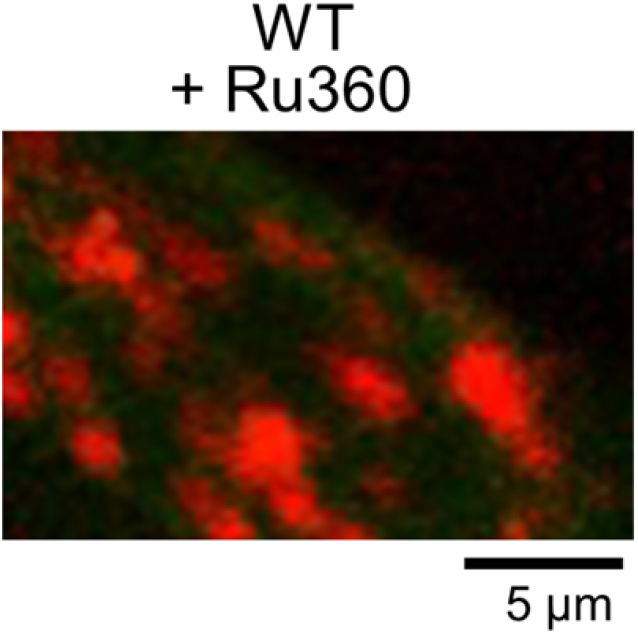
Time-lapse confocal fluorescent images of cytosolic GCaMPs ([Ca^2+^]_cyto_, green) and mtGECO ([Ca^2+^]_mito_, red) in body wall muscles of immobilized Ru360-treated WT worm (*goeIs3; aceIs1*), showing suppression of mitochondrial Ca^2+^ uptake during cytosolic Ca^2+^ fluctuations by Ru360 treatment. Images were acquired every 1 sec for 1 min. Playback, 10 fps. Relates to Fig. 3 A.

**Supporting Movie 4.**
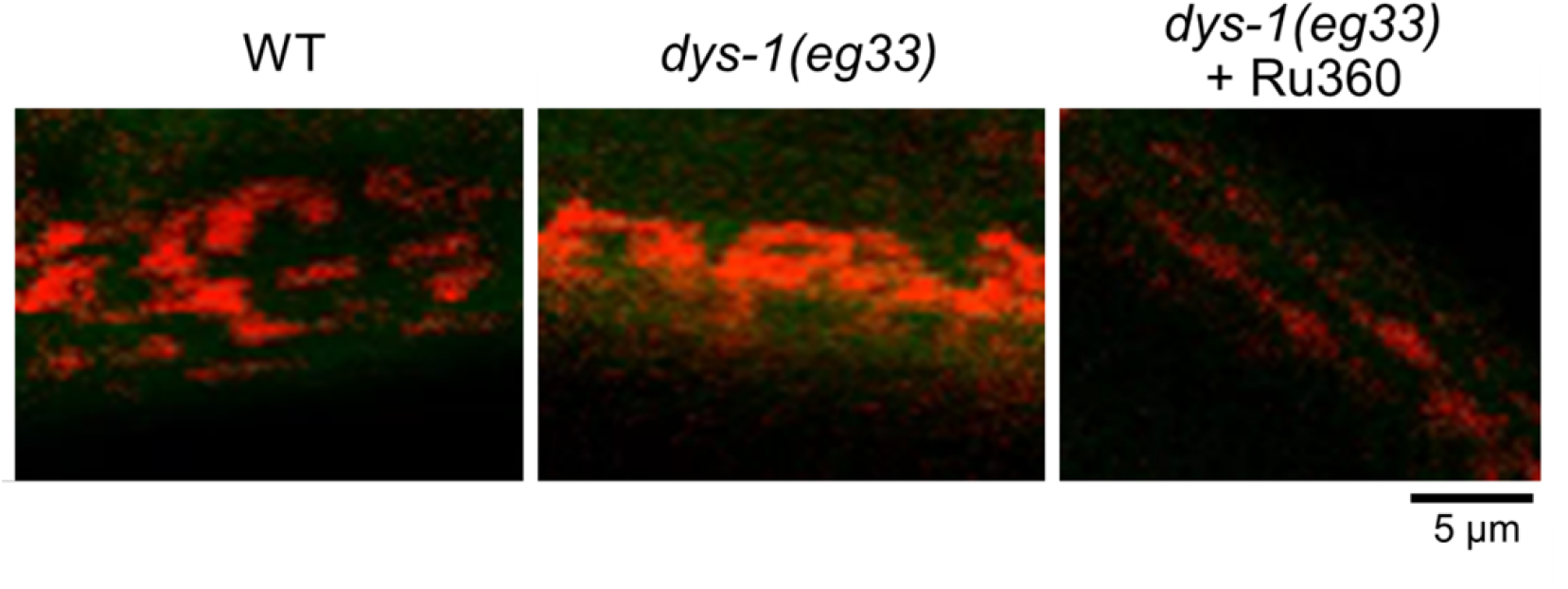
Time-lapse confocal fluorescent images of cytosolic GCaMPs ([Ca^2+^]_cyto_, green) and mtGECO ([Ca^2+^]_mito_, red) in body wall muscles of immobilized WT worm (*goeIs3; aceIs1*, left panel), *dys-1(eg33)* (center panel), and Ru360-treated *dys-1(eg33)* (right panel). Images were acquired every 1 sec for 1 min. Playback, 10 fps. Relates to Fig. 4 A.

